# Interplay Between Short-range Attraction and Long-range Repulsion Controls Reentrant Liquid Condensation of Ribonucleoprotein-RNA Complexes

**DOI:** 10.1101/600601

**Authors:** Ibraheem Alshareedah, Taranpreet Kaur, Jason Ngo, Hannah Seppala, Liz-Audrey Djomnang Kounatse, Wei Wang, Mahdi Muhammad Moosa, Priya R. Banerjee

## Abstract

In eukaryotic cells, ribonucleoproteins (RNPs) form mesoscale condensates by liquid-liquid phase separation that play essential roles in subcellular dynamic compartmentalization. The formation and dissolution of many RNP condensates are finely dependent on the RNA-to-RNP ratio, giving rise to a window-like phase separation behavior. This is commonly referred to as reentrant liquid condensation (RLC). Here, using RNP-inspired polypeptides with low-complexity RNA-binding sequences as well as the C-terminal disordered domain of the ribonucleoprotein FUS as model systems, we investigate the molecular driving forces underlying this non-monotonous phase transition. We show that an interplay between short-range cation-π attractions and long-range electrostatic forces governs the heterotypic RLC of RNP-RNA complexes. Short-range attractions, which can be encoded by both polypeptide chain primary sequence and nucleic acid base sequence, are activated by RNP-RNA condensate formation. After activation, the short-range forces regulate material properties of polypeptide-RNA condensates and subsequently oppose their reentrant dissolution. In the presence of excess RNA, a competition between short-range attraction and long-range electrostatic repulsion drives the formation of a colloid-like cluster phase. With increasing short-range attraction, the fluid dynamics of the cluster phase is arrested, leading to the formation of a colloidal gel. Our results reveal that phase behavior, supramolecular organization, and material states of RNP-RNA assemblies are controlled by a dynamic interplay between molecular interactions at different length scales.

## Introduction

Ribonucleoproteins (RNPs) form a diverse set of biomolecular condensates in eukaryotic cells by liquid-liquid phase separation (LLPS) [1]. These condensates are non-membrane bound assemblies that can dynamically exchange their components with surrounding subcellular environment [2–5]. From a physiologic point of view, RNP granules perform critical cellular functions that are conserved from yeast to humans [2, 6]. From a pathological point of view, these granules enrich many disease-linked proteins such as FUS, hnRNPA1/A2, and TDP43 [7, 8], aggregation of which is often associated with multiple degenerative disease conditions [9].

Many nuclear RNPs spontaneously self-assemble into liquid droplets *in vitro* at their physiological concentrations [10]. However, they predominantly exist in a homogeneous solution phase in cells [8, 11, 12]. This is attributed to the buffering effect of cellular RNAs [11]. Recent *in vitro* studies have demonstrated that RNAs can inhibit RNP phase separation when present at high RNA-to-RNP ratios but promote RNP phase separation at low RNA-to-RNP ratios [13]. The observed regulatory effect of RNAs is thought to be manifested via promiscuous RNA binding to evolutionarily conserved positively charged Arg/Gly-rich low-complexity domains (R/G-rich LCDs) [11, 13, 14], an RNA interacting motif present in > 50% of the eukaryotic RNPs [15, 16]. This RNA-dependent RNP condensation and subsequent decondensation can be explained by the reentrant liquid condensation (RLC) model for associative phase separation [13].

According to the RLC model for RNP-RNA phase transitions, the condensation is driven by long-range electrostatic attraction between positively charged R/G-rich disordered domains and negatively charged RNA chains. The decondensation process, however, is best explained by *charge inversion* on the surface of RNA or RNP chains. In the charge inverted state, the surface of a charged macromolecule is over-screened by oppositely charged ions, resulting in the accumulation of a greater number of counter-ions on the macromolecule than what is required to reach a charge-neutral state (Fig. 1a) [17]. With excess bound counter-ions, charge inversion triggers an energetically unfavorable long-range coulomb repulsion, which drives the progressive decondensation process. Consequently, a reentrant phase transition can be quantified using three distinct yet inter-linked parameters: (a) the condensation concentration (X_c_), (b) the decondensation concentration (X_d_), and (c) the concentration at which phase separation is maximal (X_o_) and complexes are electrostatically neutral (Fig. 1a) [17–19]. When macromolecules interact only through ionic forces, both condensation and decondensation boundaries are primarily determined by the linear charge densities and the number of charged residues in polycation (*i.e.,* R/G-rich LCD) and polyanion (*i.e.,* RNA) chains at a given buffer ionic strength [18]. Although this framework qualitatively recapitulates the reentrant nature of RNP-RNA condensation [13], its application to naturally occurring biological sequences is limited, since the RLC model ignores intermolecular short-range forces which can be sequence-specific and/or solution condition-dependent (such as molecular crowding effects [20]).

**Figure 1:**
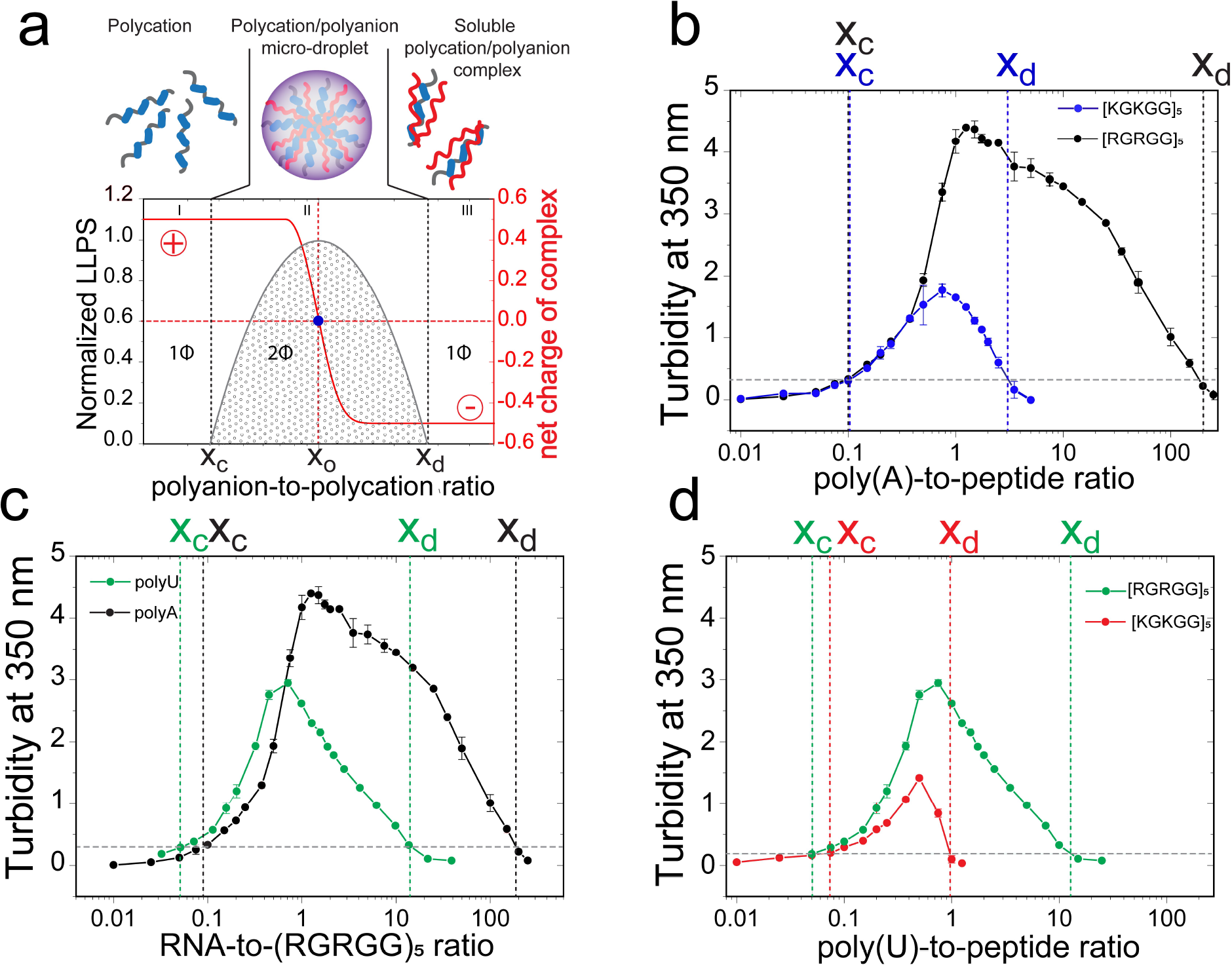
Liquid-liquid phase separation (LLPS) regimes of peptide-RNA complexes are dependent on polypeptide amino acid and RNA nucleobase sequences. **a)** A schematic representation of reentrant liquid condensation (RLC) showing various molecular species at three different polyanion:polycation regimes. Also shown here is the charge inversion of peptide-RNA complexes that results in a reentrant homogeneous phase formation at high polyanion-to-polycation ratio. **b)** Solution turbidity at 350 nm of R/G-and K/G-rich peptides and poly(A) mixtures showing that the dissolution of [RGRGG]_5_ condensates requires ~50-fold higher poly(A) concentration (black) than [KGKGG]_5_ condensates (blue). **c)** Poly(A)-[RGRGG]_5_ condensates have higher decondensation boundary compared to poly(U)-[RGRGG]_5_ condensates. **d)** [RGRGG]_5_ condensates are stable across a ~10-fold higher poly(U) concentration than [KGKGG]_5_ condensates.

Given that short-range forces are vital in controlling the RNP-RNA complexation [7, 21, 22], here we ask how sequence-encoded short-range interactions, such as the cation-π attraction, impact the reentrant liquid condensation of RNP-RNA mixtures. By measuring phase boundary curves of positively charged LCD-RNA systems and the fluid dynamics of their condensates using correlative fluorescence microscopy and optical tweezers, we show that combined intermolecular interactions on different length scales govern the phase behavior and material properties of RNP-RNA assemblies. Based on our experimental findings and theoretical modeling, we propose that long-range ionic interactions drive the phase separation of RNP-RNA complexes, while short-range attractions dictate the width of the phase separation regime as well as the viscoelastic states of the heterotypic condensates.

## Results and Discussion

### Arginine-rich and lysine-rich low-complexity sequences (LCSs) produce distinct reentrant liquid condensation behavior

Polycationic R/G-rich LCDs are prevalent in eukaryotic RNA-binding proteome [15, 16]. R/G-rich LCDs promiscuously interact with diverse RNA sequences [14, 16, 23], but their role in RNP function remains poorly understood. We and others previously reported that the R/G-rich LCDs can act as a phase separation module via multivalent long-range electrostatic interactions with RNAs [13, 14, 24]. To investigate if short-range interactions by arginine residues play any role in the phase separation behavior of R/G-rich LCD and RNA mixtures, we designed two peptide sequences: [RGRGG]_5_ and [KGKGG]_5_. Both polypeptides are estimated to carry the same overall charge (+10e at pH 7.5; http://protcalc.sourceforge.net/; Supplementary Table 1; Supplementary Fig. S3) and therefore, are expected to exhibit similar phase behavior with RNA, assuming inter-chain heterotypic ionic interactions are the only driving force. Using a homopolymeric RNA, poly(A), solution turbidity and confocal microscopy measurements revealed that both [RGRGG]_5_ and [KGKGG]_5_ peptides display reentrant liquid condensation behavior, *i.e.,* peptide-RNA droplets are formed at a low RNA-to-peptide ratio, whereas a homogeneous phase emerges at a high RNA-to-peptide ratio (Fig. 1b; Supplementary Fig. S4). The condensation boundary (x_c_) and the peak position (x_0_) were observed to be similar for both [RGRGG]_5_ and [KGKGG]_5_ peptides within the resolution of our measurements (Fig. 1b). These results are in agreement with the charge inversion mechanism [17], given the length and net charge per residue (NCPR) are the same for both peptides (Supplementary Fig. S3). However, the decondensation boundary (x_d_), which marks the re-appearance of the homogeneous phase, was ~50 fold higher for [RGRGG]_5_ when compared with [KGKGG]_5_ (Fig. 1b; Supplementary Fig. S4). Electrophoretic light scattering measurements revealed similar charge inversion transitions for both peptide systems (Supplementary Fig. S5). Repeating the phase separation measurements of [RGRGG]_5_ and [KGKGG]_5_ peptides with poly(U) RNA revealed a similar shift in the decondensation boundary, albeit the difference is less pronounced for poly(U) than poly(A) RNA (Figs. 1c&d; Supplementary Fig. S4). These data collectively suggest that despite carrying identical charges, R-rich LCDs and K-rich LCDs encode distinct reentrant phase behavior.

### Switch-like activation of short-range forces by liquid-liquid phase separation

Two key aspects of our observations on the heterotypic RNP-RNA condensation cannot be explained by a simple charge inversion mechanism [18]: (a) the decondensation boundary, but not the condensation boundary, is peptide and RNA sequence-dependent (Fig. 1b-d); (b) the experimentally determined decondensation boundaries follow a specific trend: [RGRGG]_5_-poly(A) > [RGRGG]_5_-poly(U) > [KGKGG]_5_-poly(A) > [KGKGG]_5_-poly(U) (Supplementary Fig. S6). To provide a theoretical basis for these observations, we consider a modification of the RLC model by taking peptide-RNA short-range interactions into account. *Firstly,* we consider the potential of both arginine and lysine residues to engage in cation-π interactions with an RNA base [25–30]. *Ab initio* calculations, crystal structure analyses, and molecular dynamics simulations previously established that the short-range arginine-RNA cation-π interactions are generally stronger than the lysine-RNA cation-π interactions [25, 26, 29]. Additionally, the delocalized π electrons of arginine guanidinium group can participate in π-π interactions with RNA bases [31, 32]. Finally, in addition to the electrostatic monopoles, arginine, but not lysine, contains higher order multipoles that can mediate weaker and shorter ranged directional interactions with RNA bases [32]. Therefore, arginine side chains are capable of mediating a stronger short-range multi-modal interaction network with RNA chains, as compared to lysine [32]. Indeed, our salt titration experiments reveal that ~ 3-fold higher [NaCl] is required to suppress phase separation of [RGRGG]_5_-poly(U) mixture as compared to [KGKGG]_5_-poly(U) mixture suggesting a stronger R-rich LCD-RNA interaction as compared to K-rich LCD-RNA interaction (Supplementary Fig. S7).

*Secondly*, we consider the dependence of this short-range peptide-RNA attraction on the mean intermolecular separation. While long-range electrostatic forces are operational even in a dilute polymer mixture, short-range forces are more relevant when the average inter-particle distances are small, *i.e.,* at relatively high volume fractions of the peptide and RNA [33–36]. Since the molecular packing density within a phase separated liquid droplet (*regime-II* in Fig. 1a) is manifold higher than a dilute peptide-RNA mixture (*regime-I* in Fig. 1a) [37] (Supplementary Fig. S8), we propose that the short-range forces are *activated* via peptide-RNA condensation, which is driven by long-range electrostatic interactions.

Based on these considerations, we formulate a modified potential to describe the reentrant phase behavior of RNP-RNA mixtures that contains a short-range attractive term activated at the condensation boundary (Fig. 2a; Supplementary Note-1) and added to the charge inversion potential [17]. The charge inversion potential accounts for both electrostatic attraction and repulsion as determined by the RNA-to-RNP ratio [18]. At the charge-neutral point, the electrostatic repulsion is minimized due to equal number of charges on each complex, and phase separation is favored. Past the charge-neutral point, the net charge on each complex is negative (Supplementary Fig. S5), resulting in a long-range coulomb repulsion that progressively destabilizes the condensates with increasing RNA concentration. However, in presence of a short-range attraction, the condensates are stabilized against the ionic repulsive force. Therefore, a competition between the long-range repulsion and short-range attraction results in effective widening of the phase separation window due to the decondensation boundary being shifted to higher RNA concentrations without any significant alterations in the onset of condensation (Fig. 2b).

**Figure 2:**
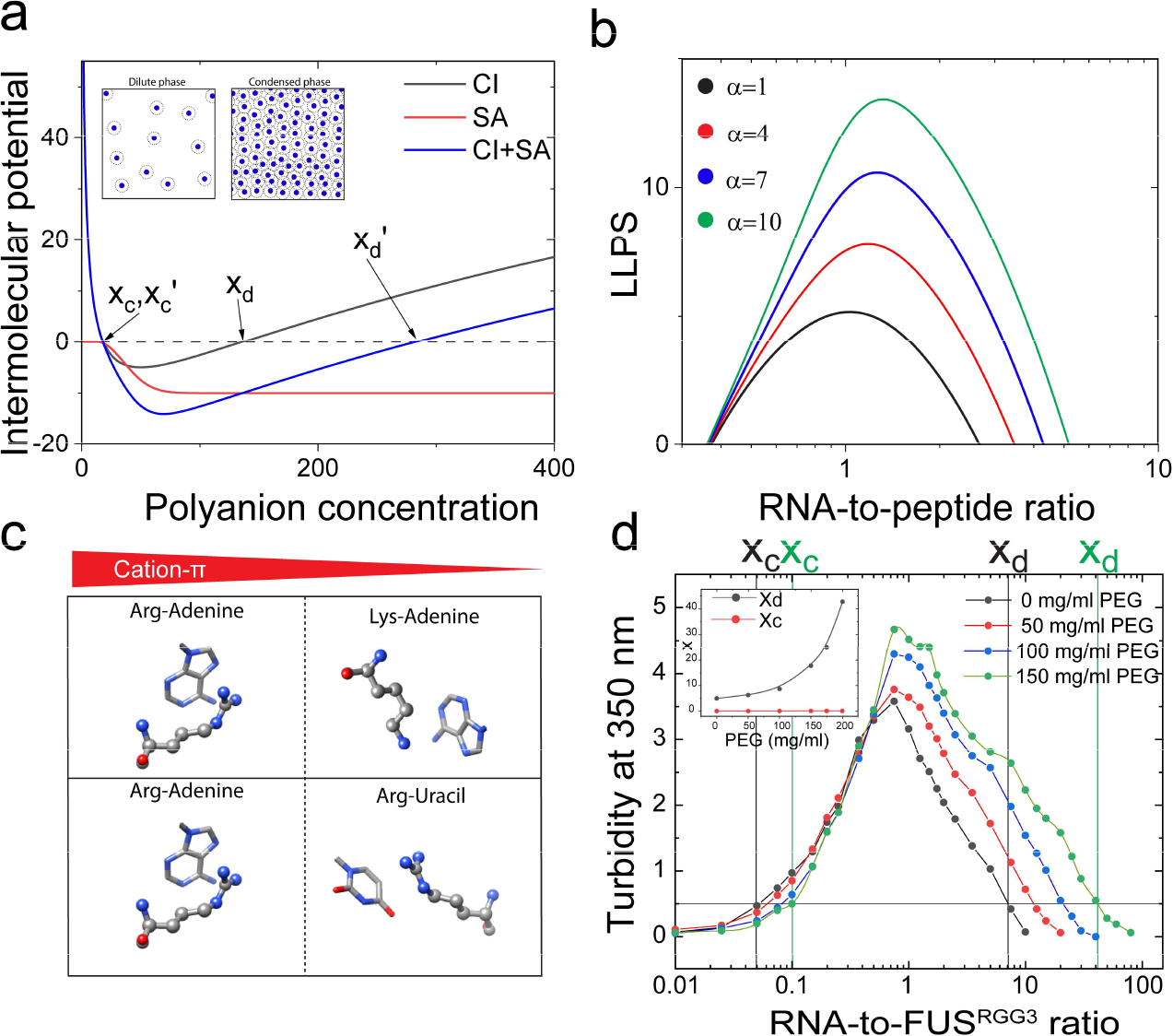
Short-range attraction is activated by the LLPS of Arg-rich peptide-RNA complexes. **a)** Molecular pair potential diagram representing <u>c</u>harge inversion potential (CI; blue) and a generalized <u>s</u>hort-range <u>a</u>ttractive potential that is activated upon condensation (SA; red). The *insets* show a schematic highlighting that the mean inter-molecular distance is considerably reduced upon phase separation, which drives the switch-like activation of the short-range forces. **b)** Simulated phase boundary curves with increasing short-range attractions (parameter α is the depth of the step function, SA in a). **c)** Schematics showing cation-π interactions between Arg/Lys-nucleobases and their relative strengths. **d)** Solution turbidity data showing an increase in the decondensation boundary upon increasing crowder concentration in FUS^RGG3^-poly(U) mixture (FUS^RGG3^: 0.33 mg/ml = 100 µM, buffer: 25 mM Tris-HCl, pH 7.5, 20 mM DTT, 150 mM NaCl). *Inset*: decondensation boundary increases with PEG8000 (x_d_; black) while condensation boundary remains relatively unaltered (x_c_; red).

Next, utilizing our model, we evaluate how the RLC behavior is affected upon variations in the strength of short-range peptide-RNA attractions. Simulated phase boundary curves suggest that the decondensation boundary monotonically moves towards higher RNA-to-peptide ratios with increased short-range attraction (Fig. 2b). We postulate that our experimentally observed trend in peptide-RNA decondensation boundaries (Supplementary Fig. S6) is correlated with the strengths of the short-range interactions between positively charged amino acids and π electron systems of nucleobases.

The exact order of cation-π interaction strengths in protein-RNA complexes are yet to be determined experimentally. However, previous computational studies provide some insights into R/K-nucleobase interactions [25–30]. Firstly, as discussed before, from the peptide sequence perspective, R-rich LCDs have more pronounced short-range attraction as compared with K-rich LCDs. Secondly, from the ribonucleic acid sequence perspective, purine bases are more efficient at forming cation-π bonds as compared with pyrimidine bases [25, 26, 29] (Fig. 2c). Therefore, [RGRGG]_5_-poly(A) cation-π interactions are expected to be the most favorable and [KGKGG]_5_-poly(U) interactions will be the least favorable amongst our four possible peptide-RNA combinations. Quite interestingly, we observe that [RGRGG]_5_-poly(A) condensates resist the reentrant dissolution the most, whereas [KGKGG]_5_-poly(U) condensates resist the dissolution the least (Fig. 1b-d; Supplementary Fig. S6). This suggests that the observed rank ordering of the RNP-RNA condensate dissolution is indeed due to the variable strengths of R/K-nucleobase interactions.

Next, we hypothesized that if short-range cation-π interactions between peptide arginine and RNA nucleobases are primarily responsible for the observed differences between the RLC of R-rich and K-rich LCDs, then abolishing the cation-π interactions by removing nucleobases from the RNA chain should result in superimposed RLC curves for the two LCD types. To test this, we characterized the RLC behavior of R-rich and K-rich LCDs upon their interactions with polyphosphate (a model for the negatively charged backbone of nucleic acids that lacks nucleobases). As expected, both [RGRGG]_5_ and [KGKGG]_5_ peptides displayed identical phase separation windows with polyphosphate, supporting the idea that cation-π interactions are primarily responsible for the observed differences in cationic LCD phase behavior with RNAs (Supplementary Fig. S9).

Finally, to test whether the observed effects are generic to short-range forces and are not specific to cation-π interactions, we introduced a tunable short-range attraction between peptide and RNA chains using a polymer crowder. Previous reports [37–39] revealed that molecular crowding by “inert” non-ionic polymers, such as polyethylene glycol (PEG), in protein/RNA-buffer solutions can introduce isotropic short-range intermolecular attraction via the depletion effect [40]. The range and strength of PEG-mediated short-range attraction are controlled by the molecular weight and concentration of PEG chains, and are typically modelled by an adjustable Lennard-Jones potential [41]. We determined phase boundary curves of mixtures of a naturally occurring R/G-rich LCD motif of FUS (FUS^RGG3^; Fig. 2d; Supplementary Fig. S3c) with poly(U) RNA at various PEG8000 concentrations. With increasing PEG from 0 to 150 mg/ml, the condensation boundary (x_c_) remained unaltered for FUS^RGG3^-poly(U) mixtures, whereas the decondensation boundary (x_d_) shifted to higher RNA-to-FUS^RGG3^ ratio by almost an order of magnitude (Fig. 2d). Therefore, switch-like activation of any short-range attraction within the peptide-RNA liquid condensates extends the phase separation regime over a broader range of RNA-to-polypeptide concentrations. Taken together, our experimental data combined with the proposed model provide a mechanistic picture on RNP-RNA RLC — long-range electrostatic attraction drives associative peptide-RNA liquid condensation in a dilute solution, whereas a competition between short-range attraction and long-range electrostatic repulsion controls the decondensation phase boundary.

### Synergy between short-range and long-range forces controls the material properties of peptide-RNA condensates prior to a charge inversion

The material properties of complex fluids are collectively determined by the net strength of respective intermolecular interaction networks [42–44]. The combination of a tunable short-range force and a non-monotonic long-range electrostatic interaction between polypeptides and RNA chains (Figs. 1a&2a) provides a unique potential for controlling the material properties of these condensates. We hypothesized that this can be achieved by a systematic variation of polypeptide and RNA sequence as well as their mixture compositions, which tunes short-range and long-range ionic forces, respectively. Before and at the charge neutral state (≤ x_0_), our model predicts that short-range cation-π interactions act synergistically with electrostatic attraction. Assuming the long-range ion-ion interactions are similar in both K/G-rich and R/G-rich peptide systems with a given RNA sequence (Supplementary Fig. S3), we hypothesized that the material properties of peptide-RNA condensates will be determined by the net strength of short-range attraction between respective polypeptide and RNA chains. Based on our experimentally determined phase boundary curves (Fig. 1b), we anticipate that R/G-poly(A) droplets will display slower fluid dynamics than K/G-poly(A) droplets. To test this idea, we employed two complementary techniques: Fluorescence Recovery After Photobleaching (FRAP) and controlled fusion of suspended peptide-RNA droplets using a dual-trap optical tweezer. Our FRAP experiments reveal that peptide diffusion is nearly arrested within R/G-poly(A) droplets as compared to the peptide diffusion within K/G-poly(A) condensates (Fig. 3a & Supplementary Fig. S10). Furthermore, a greater degree of fluorescence recovery was observed for R/G-poly(U) droplets as compared to R/G-poly(A) droplets. These data are consistent with the idea that *in-droplet* peptide mobility at the nanoscale is dependent on the inter-molecular friction due to short-range interactions (cation-π forces in our case) with RNA, and with increasing strengths of such interactions *in-droplet* diffusivity dynamics become slower [45]. The observed trend in FRAP results is also in agreement with the rank order of the energetics of short-range interactions of amino acid-base combinations [25, 26, 29].

**Figure 3:**
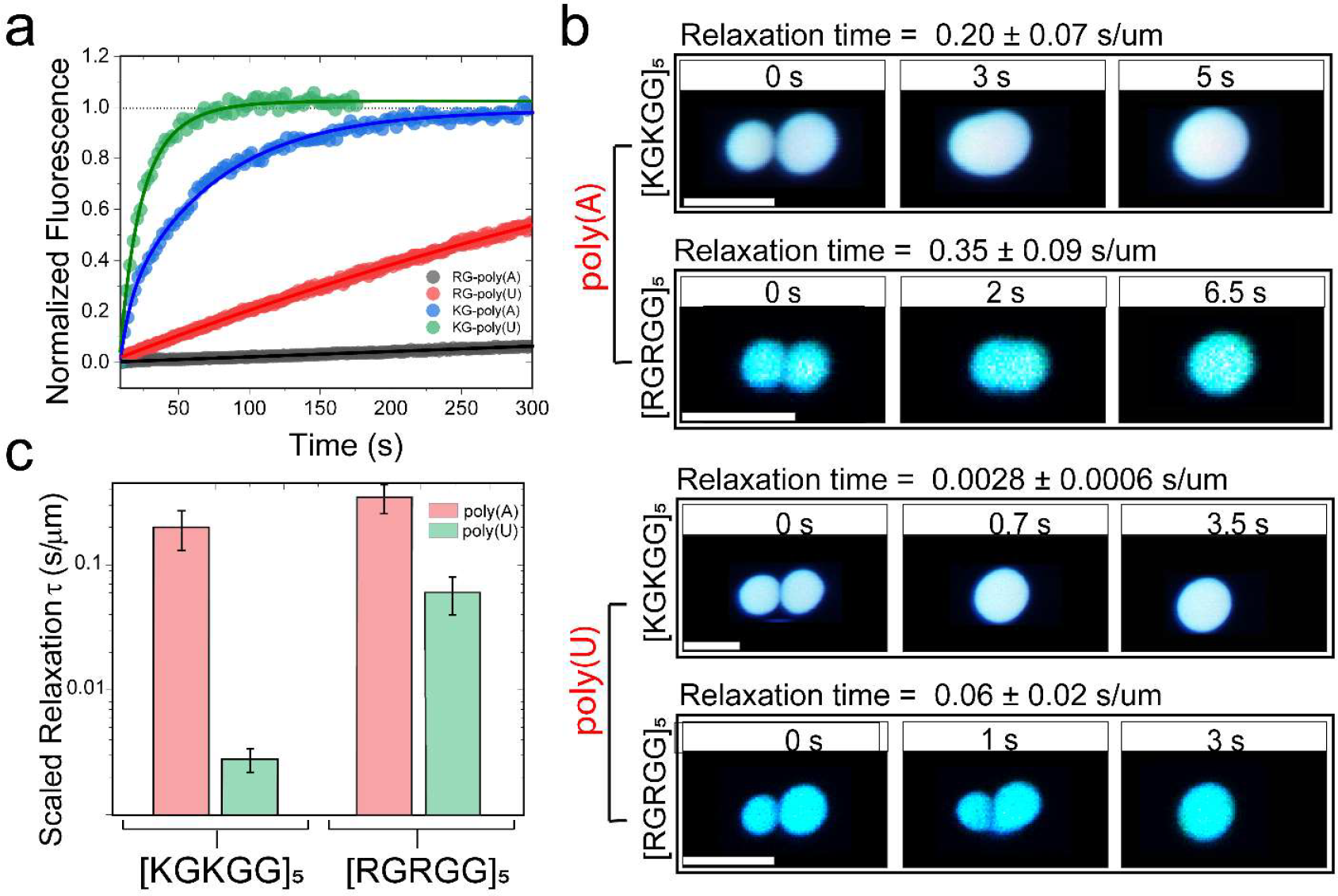
Short-range attraction tunes material states of RNA-peptide condensates. **a)** Normalized fluorescence recovery after photobleaching (FRAP) intensity time-trace for peptide/RNA systems. The mobile phase fraction (ratio of final intensity to pre-bleach intensity) and recovery time show a scaling with the strength of the short-range attraction between RNA and the peptide. R/G-poly(A) (black): no recovery observed with < 5% mobile fraction. KG-poly(U) (green): fastest recovery with ~100% mobile fraction. **b)** Time-lapse images of trap-induced fusion of RNA-[RGRGG]_5_ droplets showing higher fusion relaxation time as compared to RNA-[KGKGG]_5_ droplets for both poly(U) and poly(A) RNA. Corresponding movies are provided as Supplementary Movies 1-4. **c)** Relaxation time of the studied peptide-RNA condensates with poly(U) (green) and poly(A) (red). [KGKGG]_5_-poly(U) condensates display the fastest relaxation time, whereas [RGRGG]_5_-poly(A) condensates display the slowest relaxation time. Scale bars in **b** represent 5 µm. All samples were prepared at 1.0 mg/ml peptide concentration, 0.75 mg/ml RNA concentration (corresponding to the peak in their turbidity plots shown in Fig. 1b-d) in 25 mM Tris-HCl, 20 mM DTT and 25 mM NaCl (pH = 7.5).

Simultaneously, to probe for the dynamics of peptide-RNA condensates at the micron-scale, we performed force-induced droplet coalescence using an optical tweezer system (Figs. 3b&c). In these experiments, two droplets were optically trapped and brought slowly into contact to initiate coalescence. The fusion timescale, which provides direct insights into the material state of the condensates [46], is estimated from the force relaxation curves measured by the traps operating at ~12 µs time resolution. Similar to our observation in FRAP experiments, K/G-poly(U) droplets displayed the fastest relaxation time of coalescence (0.0028±0.0006 s/µm), which indicates a high degree of fluidity and low viscosity (Figs. 3a&b), whereas R/G-poly(A) droplets showed more than two orders of magnitude slower relaxation time (Fig. 3c). In a similar vein, a progressive increase in droplet fusion time was confirmed for FUS^RGG3^-poly(U) droplets with increasing depletion attraction by PEG (Supplementary Fig. S11). Together with the FRAP results, the fusion data confirms that a combination of short-range attraction and long-range forces controls the molecular diffusion and the mesoscale fluid dynamics of peptide-RNA condensates. We note that similar observations on the material properties of PR protein-RNA mixtures were recently reported by Boeynaems *et al.*[31], which are in agreement with our results reported here.

### RNA sequence tunes material states of FUS^R/G-rich LCD^ condensates

Our results using RNP-inspired polypeptide sequences indicate that the dynamics of the phase separated condensates formed by R/G-rich LCDs are sensitively dependent on RNA-base sequences. We next tested whether this effect is extended to naturally occurring RNA binding proteins with complex sequence patterns. Using an RNA binding domain construct of FUS [FUS^R/G-rich LCD^; contains three disordered R/G-rich LCDs and one folded RNA recognition motif (RRM)] (Supplementary Fig. S3d), we employed FRAP and trap-induced fusion experiments to investigate the mesoscale RNP-RNA condensate dynamics (Fig. 4a). In presence of poly(A) RNA, FUS^R/G-rich LCD^ condensates showed almost an order of magnitude slower fusion kinetics than condensates formed by poly(U) RNA. Simultaneously, FRAP measurements revealed ~ 2-fold decrease in protein diffusion within poly(A)-FUS^R/G-rich LCD^ condensates as compared to the poly(U)-FUS^R/G-rich LCD^ droplets (Fig. 4b). Similar behavior was also observed for the isolated third RGG box of FUS (FUS^RGG3 (472-505)^; Supplementary Fig. S3) with more pronounced difference in fusion relaxation timescales for FUS^RGG3^ condensates with poly(A) and poly(U) RNAs (Fig. 4c&d). These results suggest that RNA base sequence tunes the material states of RNP-RNA condensates through short-range cation-π interactions.

**Figure 4:**
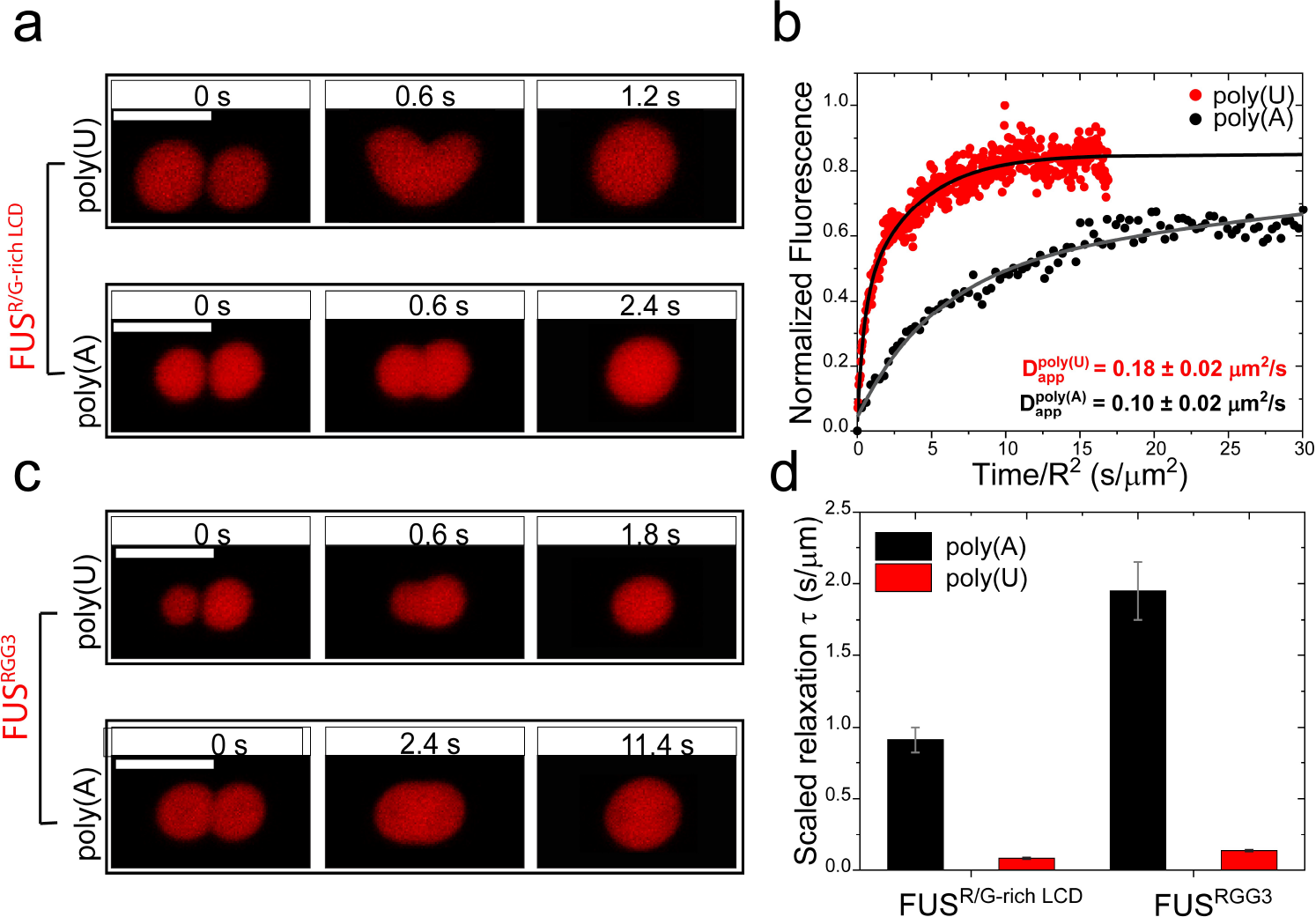
RNA base sequence tunes material states of FUS RNA-binding domain condensates. **a)** Trap-induced fusion time lapse images of poly(U) and poly(A) RNA with FUS^R/G-rich LCD^. **b)** Normalized Fluorescence recovery after photobleaching (FRAP) intensity time-trace for FUS^R/G-rich LCD^-RNA condensates. The recovery time (x-axis) was normalized with respect to the radius of the bleaching ROI to account for the differences in the bleaching regions of interests (ROIs). **c)** Trap-induced fusion time lapse images of poly(U) and poly(A) RNA with FUS^RGG3^. **d)** Relaxation times of the FUS-RNA condensates are approximately one order of magnitude higher for poly(A) as compared to poly(U). Scale bars in a and c represent 5 µm. Corresponding movies for a and c are provided as Supplementary Movies 5 and 6.

### Competition between short-range and long-range forces leads to the emergence of a cluster phase at a high RNA-to-RNP ratio

Reentrant liquid condensation of protein-RNA complexes is manifested by a highly non-monotonic electrostatic potential, which is sensitively dependent on the RNA-to-polypeptide ratio (Fig. 1a). So far, studies pertaining to electrostatically-driven biological phase separating systems are mostly limited to mixture compositions corresponding to the charge-neutral state (x_0_), and hence, little is known about the structure and material state of the condensates past the charge inversion point (*i.e.,* at RNA-to-peptide ratios greater than respective x_0_’s) where a long-range repulsive force operates. In contrary to the charge neutral state for our peptide-RNA condensates, confocal microscopy of peptide-RNA mixtures past respective x_0_ points reveals the emergence of a *cluster phase* where small spherical peptide-RNA assemblies aggregated together to form a colloid-like solution that did not coalesce (Fig. 5a; Supplementary Movie 7). This behavior is more pronounced at lower buffer ionic strength (≤ 25 mM Na^+^) where electrostatic interactions exist over a longer range (i.e., higher Debye length). Using electrophoretic light scattering measurements, we confirmed that these condensates are strongly negatively charged (Supplementary Fig. S5), suggesting that the suspension of colloid-like condensates is kinetically stabilized by long-range electrostatic repulsions. This scenario is analogous to colloidal clusters formed by micro-particle suspensions where the colloidal particles interact via a mixed attractive and repulsive potential [47]. Under such conditions, combining long-range repulsion with a weaker short-range attraction between colloidal particles in a suspension results in the formation of a dynamic cluster phase, while increasing inter-particle attraction leads to colloidal gel formation with cluster-like morphologies [36, 47, 48]. Remarkably, we observed that while R/G-poly(U) droplet clusters displayed relatively fast diffusion dynamics (Fig. 5a), R/G-poly(A) droplet clusters showed no FRAP recovery under similar conditions (Fig. 5b). These data suggest a severely restricted *in-droplet* fluid dynamics for R/G-poly(A) clusters, which is consistent with the stronger Arg-adenine interactions as compared to the Arg-uracil pair. To investigate whether R/G-poly(A) condensates exchange their components within a single cluster, we mixed independently prepared R/G-poly(A) clusters of different colors (using independently labeled peptides with Alexa488 and Alexa594 dyes). Two-color confocal microscopy imaging revealed that R/G-poly(A) clusters indeed can coexist in a complex mixture and retain the internal composition for an extended period of time (Fig. 5c). Contrastingly, R/G-poly(U) clusters exchange their materials rapidly under identical conditions (Fig. 5d). Weakening the electrostatic interactions by increasing the ionic strength of the buffer, we observed that both R/G-poly(A) and R/G-poly(U) clusters transitioned to a bulk liquid phase by coalescence (Supplementary Fig. S12). Interestingly, cluster phase was not observed for K/G-poly(A) mixtures under similar conditions, and instead a relatively homogeneous suspension of negatively charged spherical condensates was visible (Supplementary Fig. S13). These results suggest that short-range forces are a critical determinant for the colloid-like cluster formation in peptide-RNA mixtures. Therefore, we conclude that a competition of sequence encoded short-range attraction and long-range repulsion amongst peptide-RNA complexes can give rise to distinct supramolecular organization of peptide-RNA condensates at higher length-scales.

**Figure 5:**
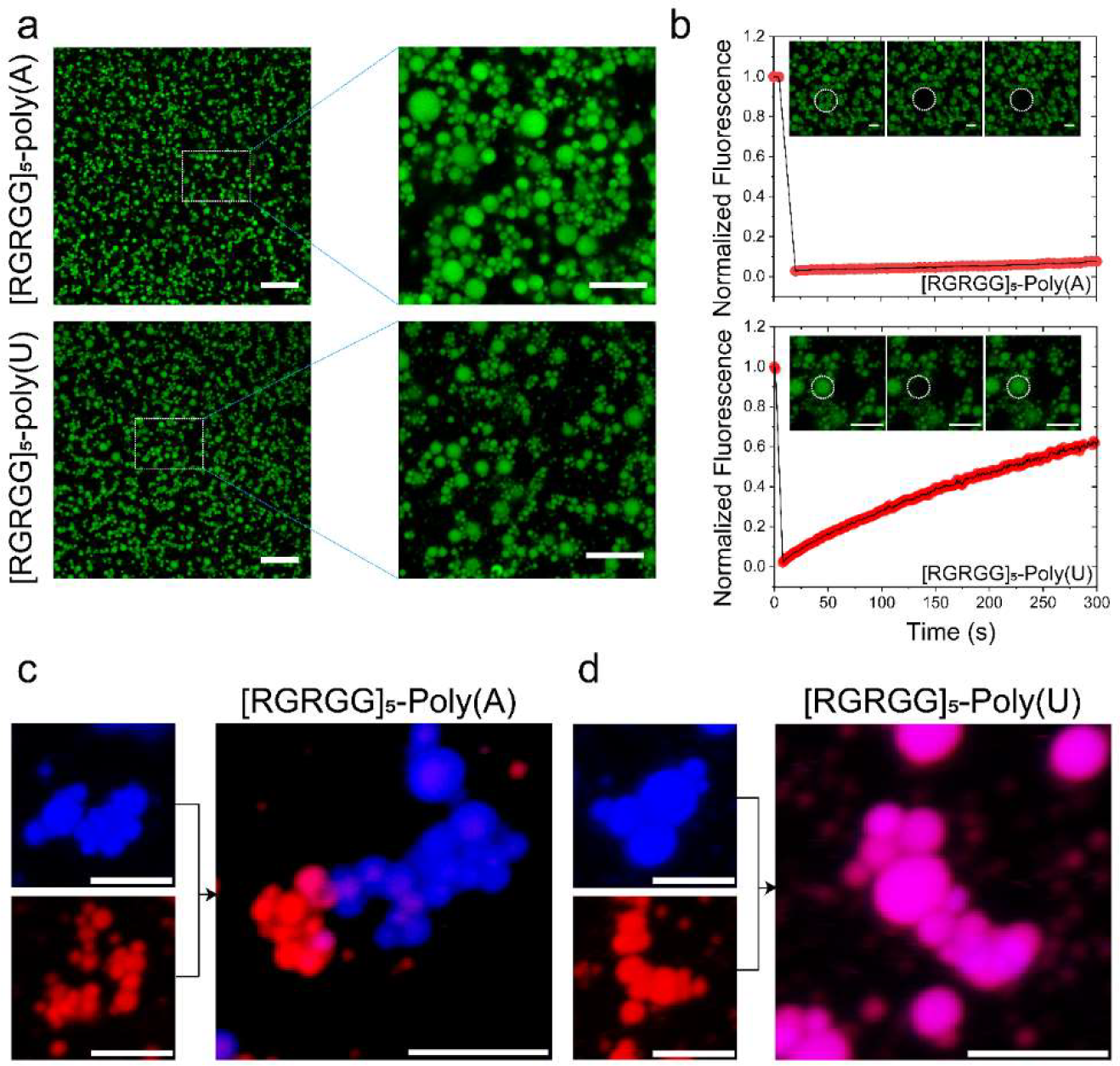
Long-range ionic repulsion competes with short-range attraction at high RNA-to-peptide ratio leading to a colloid-like cluster phase: **a)** Fluorescence micrographs showing cluster formation for [RGRGG]_5_-poly(A) *(top)* and [RGRGG]_5_-poly(U) condensates *(bottom)*. Scale bar represents 20 µm (left) and 10 µm (right). **b)** RNA sequence tunes the fluid dynamics of [RGRGG]_5_ clusters showing arrested diffusion in poly(A) clusters and a more dynamic diffusion in poly(U) clusters as evidenced by the fluorescence recovery time traces. Scale bars represent 5 µm for poly(A) images and 4 µm for poly(U) images. Corresponding movies are provided as Supplementary Movies 8 and 9. **c)** [RGRGG]_5_-poly(A) condensates can form co-clusters that retain their composition in a complex mixture. **d)** The [RGRGG]_5_-poly(U) condensates exchange their content rapidly. Scale bar represents 5 µm for both **(c)** and **(d)**.

### Conclusion

Electrostatic forces play a fundamental role in driving the formation of biomolecular condensates in the cell [49]. Previously, we reported reentrant liquid condensation behavior of R/G-rich LCDs with RNA [13], a phenomenon that can explain the enhanced solubility of prion-like RNA binding proteins in the nucleus (high RNA environment) [11]. In this work, we explore how reentrant phase transition is controlled by varying RNA and polypeptide sequences at the molecular level. We demonstrate that intermolecular long-range ionic interactions act as a “steering force” that drives liquid condensation of peptide/RNA complexes in a dilute mixture. Upon the formation of a phase separated complex (condensate), short-range forces become significant due to the increased *in-droplet* intermolecular proximity. These short-range forces stabilize phase separated complexes against reentrant dissolution at higher RNA-to-polypeptide ratios. Sequence-encoded short-range forces, such as cation-π interactions, enable fine control of the width of the phase separation window. Combined with the long-range ionic forces, these sequence-specific interactions can collectively determine the material states of polypeptide-RNA condensates. Although our current study focuses on the cationic LCD and RNA mixtures, the physical principles presented here are generally applicable to other biological systems (*e.g.,* Alzheimer protein Tau [50]) where phase separation is electrostatically driven. Additionally, we envision that posttranslational modifications that alter physicochemical properties of cationic amino acid residues can tune the phase behavior and fluid properties of cellular RNP condensates.

Unlike short-range attractions, electrostatic interactions (attractive *vs*. repulsive) are non-monotonic and are determined by the overall stoichiometry of respective complexes (Fig. 1a). This offers an intuitive means of tuning the structure and dynamics of the RNA-protein condensates by varying the heterotypic mixture composition. When complexes are neutral, the short-range and long-range forces act synergistically to determine the material properties of RNP-RNA condensates (Figs. 3&4). At higher RNA-to-peptide ratios (where complexes are negatively charged), a competition between short-range and long-range forces can lead to the formation of a colloid-like phase, which is marked by the formation of droplet clusters. Using peptide (R or K) and RNA sequence (purine or pyrimidine) variants, we show that the structure and dynamics of these clusters are also dependent on the strength of cation-π interactions. With relatively high short-range attraction between the R/G-rich peptide and poly(A) chains, these clusters behave analogously to a colloidal network gel that can be formed via a mixed interaction potential [41, 47]. These results indicate that the rules of sequence-specific molecular interactions are relevant not only to understand the reentrant liquid condensation behavior of RNP-RNA complexes, but also to potentially engineer new mesoscale materials where viscoelastic peptide-nucleic acid microspheres could be utilized as soft colloids for gelation. Unlike commonly used DNA and protein-coated polystyrene particles [51, 52], the hardness of these peptide-RNA microspheres can be finely tuned by varying amino acid and nucleobase sequences of polypeptides and nucleic acids, respectively.

## Acknowledgements

The authors gratefully acknowledge UB north campus confocal imaging facility (supported by National Science Foundation MRI Grant: DBI 0923133) and its director, Mr. Alan Siegel for helpful assistance.

## Funding

We gratefully acknowledge support for this work from University at Buffalo, SUNY, College of Arts and Sciences to P.R.B.

## Author contributions

P.R.B. designed the study. I.A., T.K., and P.R.B. designed the experimental strategies with input from M.M.M. W.W. and T.K. expressed and purified recombinant proteins from bacteria. T.K., W.W., L-A.D.K., and J.N. performed the phase diagram analysis. I.A. collected confocal microscopy experiments, partitioning, FRAP measurements, and trap-induced droplet fusion with help from P.R.B. and T.K. I.A. and P.R.B. performed the phase separation model building and numerical simulation. P.R.B., I.A., T.K., and M.M.M. wrote the manuscript.

## Conflicts of Interest

The authors declare no conflict of interest.

## Supporting Information

Contains Materials and Methods; Supplementary Figures S1-S13; Supplementary Movies 1-9; Supplementary Table-1; Supplementary Note-1; Supplementary References.

