## Supplementary File 1 for "Interplay Between Short-range Attraction and Long-range Repulsion Controls Reentrant Liquid Condensation of Ribonucleoprotein-RNA Complexes"

### <u>Supplementary Information</u>

**Materials and Methods**:

*Peptide and RNA sample preparation:* $[RGRGG]_5$, $[KGKGG]_5$, and RGG-3 domain of FUS ($FUS^{RGG3}$:472-505) were synthesized by GenScript USA Inc. (NJ, USA; ≥ 95% purity) and were used without further purification. All peptide sequences contain a C-terminal cysteine for site specific fluorescence labeling. Concentrated stock solutions were made using RNase-free water (Santa Cruz Biotechnology) containing 50 mM dithiothreitol (DTT) to prevent cysteine oxidation. Polyuridylic acid (poly(U); molecular weight = 600-1000 kDa) and polyadenylic acid (poly(A); molecular weight = 100-500 kDa) were purchased from Sigma-Aldrich. RNA stock solutions were prepared in RNase-free water (Santa Cruz Biotechnology) and concentrations were determined by the absorbance at 260 nm using a NanoDrop oneC UV-Vis spectrophotometer. Inorganic polyphosphate (poly(P); medium chain; ~ 45-160 phosphate units) was purchased from Kerafast (Boston, MA) and stock solutions were made using RNase-free water (Santa Cruz Biotechnology).

*Protein samples:* The C-terminal RNA-binding domain of FUS ($FUS^{R/G\text{-}rich\ LCD}$: 211-526Δ422-453) was prepared as described before[1]. $FUS^{R/G\text{-}rich\ LCD}$ was expressed in E. coli cells (BL21(DE3)) and then extracted using a french press. $His_6$-tagged proteins were subsequently purified using Ni-NTA agarose matrix. To check the purity of the proteins, polyacrylamide gel electrophoresis (PAGE) and Coomassie blue staining were used. Concentrations of the protein samples were determined using absorbance at 280 nm (extinction coefficient: 86,750 $M^{-1}.cm^{-1}$ for His6-MBP-$FUS^{R/G\text{-}rich\ LCD}$; https://web.expasy.org/protparam). The protein samples were flash frozen in individual aliquots, stored at -80 °C and thawed prior to experiments.

*Fluorescence labeling of peptides and proteins:* Individual peptides and the A313C variant of $FUS^{R/G\text{-}rich\ LCD}$ were site-specifically labelled using Alexa488 or Alexa594 dyes (C5-maleimide derivative, Molecular Probes) using cys-maleimide chemistry as described in our earlier works[1-5]. Briefly, the labelling reactions were carried out at 4 °C overnight in the dark. Excess free dyes from $FUS^{R/G\text{-}rich\ LCD}$ reaction mixture were removed by centrifugal filtration with a 3K cutoff filter (Millipore). For the peptides, excess dyes were removed by four rounds of acetone precipitation. Four times the sample volume of cold (-20°C) acetone was added to the reaction mixture. The reaction was then vortexed and incubated at -20°C for 1-2 hrs, followed by centrifugation at 12,000 x g for 10 minutes and decantation of the supernatant. After four rounds, the acetone was allowed to evaporate and the resulting dry purified labelled peptide pellet was resuspended in RNase-free water. Purities of the labelled proteins and peptides were tested via SDS-PAGE and mass spectrometry. A313C variant of $FUS^{R/G\text{-}rich\ LCD}$ was expressed and purified using similar protocol as the wild-type protein. UV-Vis absorption measurements were used to measure the labelling efficiencies (≥ 85% in all cases).

*Solution Turbidity measurements:* Peptide and RNA mixtures were prepared at 100 μM peptide concentration with variable RNA concentrations. The buffer contained 25 mM Tris-HCl (pH 7.5) and 20 mM DTT. The absorbance was measured at 350 nm using a NanoDrop oneC UV-Vis spectrophotometer at room temperature after ~100 seconds of sample equilibration with a 1.0 mm optical path length. Each turbidity plot was generated via gradual RNA titration[3]. Measurements were performed in triplicates. The phase boundary curve was obtained by plotting the data using OriginPro software. For the crowding experiments, the experimental buffer contained desired amounts of polyethylene glycol (PEG8000; Sigma-Aldrich) as reported in respective figure legends. The titrations were carried out in a similar manner.

*Phase diagram analysis:* Binary phase diagrams were constructed by measuring phase boundary curves of peptide-RNA mixtures with variable starting concentrations of a given peptide using turbidity measurements in conjunction with optical microscopy. Each sample was first subjected to turbidity measurement and subsequently placed under a Primo-vert inverted iLED microscope (Zeiss; using either 40x or 100x objective lens), equipped with a Zeiss Axiocam 503 monochorme camera. The droplets were clearly visible for samples with a measured solution turbidity value of ~ 0.5 or higher (10 mm path length) at 350 nm.

*Fusion of suspended droplets using optical traps:* Controlled fusion assays were conducted to investigate the mesoscale dynamics of RNA-peptide droplets, as previously described[1]. Briefly, each sample was injected into a tween 20-coated (20% v/v) 25 mm x 75 mm x 0.1 mm single chamber of the custom-made flow cell. Samples were prepared at concentrations that correspond to the peak in solution turbidity plot, i.e., 1.00 mg/ml peptide and 0.75 mg/ml RNA, in a buffer containing 25 mM Tris-HCl, 25 mM NaCl, and 20 mM DTT (pH 7.5). For $FUS^{RGG3}$ fusion experiments, samples were prepared at 0.33 mg/ml peptide concentration and 0.25 mg/ml RNA concentration (RNA:peptide = 0.75) in a buffer containing 25 mM Tris-HCl, 125 mM NaCl, and 20 mM DTT (pH 7.5). $FUS^{R/G\text{-rich LCD}}$ samples were prepared at 1.0 mg/ml protein and 0.075 mg/ml RNA in the same buffer as $FUS^{RGG3}$. Droplets were trapped using a dual-trap optical tweezer system coupled to a laser scanning confocal fluorescence microscope (LUMICKS™, C-trap). Two droplets were independently trapped using optical traps operating at minimum power (to minimize heating the trapped droplets) and then brought into close proximity. The trapping of the droplets by the 1064 nm laser was achieved via the significant difference in the refractive index between the droplet and dispersed phases. Trap-2 was kept fixed in space and trap-1 was set to move at a constant speed of 100 nm/s in the direction of trap-2. Once the trapped droplets fuse and relax to a spherical shape, the trap motion was stopped and the force-time signal was analyzed. The force on the moving trap (i.e., trap-1) was recorded at 78.4 kHz sampling frequency (i.e., ~12 μs time interval) and analyzed using an appropriate fusion relaxation model[6]. The following equation was used to fit the force-time curve:

$$F = ae^{(-t/\tau)} + bt + c \tag{1}$$

Where the parameter $\tau$ is the fusion relaxation time. The 2nd term in equation (1) is used to account for the trap's constant velocity. We recorded at least 10-15 controlled fusion events per sample, then scaled every relaxation time by the average size of the fusing droplets. A representative force relaxation curve is shown in Figure S1.

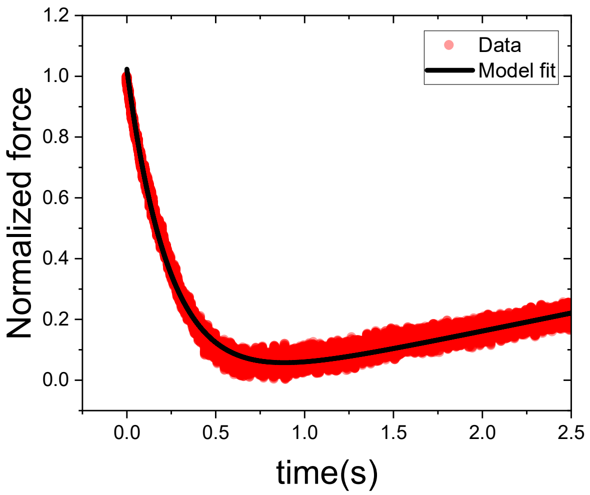


**Figure S1:** Representative normalized force relaxation curve during trap-induced coalescence of suspended peptide-RNA droplets. The data (red trace) is fitted (black line) using a previously published model described by the equation-1.

*FRAP measurements and confocal images of the condensates:* Fluorescence recovery after photobleaching (FRAP) was measured using a Zeiss LSM710 laser scanning confocal microscope, equipped with a 63x oil-immersion objective (Plan-Apochromat 63x/1.4 oil DIC M27). Samples were incubated in a tween-coated (20% v/v) Nunc Lab-Tek Chambered Coverglass (ThermoFisher Scientific Inc.) at room temperature. For FRAP measurements, samples were prepared by mixing 1.00 mg/ml of peptide and 0.75 mg/ml of RNA (corresponding to the peak position in their turbidity plot) in 25 mM Tris-HCl, 20 mM DTT, and 25 mM NaCl (pH 7.5). Alexa488-labeled peptides (excitation/emission wavelengths were 488 nm/503-549 nm) were used as fluorescence probes in these experiments. Each FRAP curve was obtained by bleaching a specific region of interrogation (ROI) for ~ 6 s with the maximum available laser power and subsequently recording the intensity trace of the bleached ROI for approximately 300 seconds or until full recovery. FRAP curves were corrected for photo fading using a reference ROI from an unbleached droplet. Normalized and corrected intensity time-traces were plotted using OriginPro software and corresponding images were processed using Fiji[7]. For FRAP measurements of the $FUS^{R/G\text{-}rich\ LCD}$ samples, a protein concentration of 1.0 mg/ml and an RNA concentration of 0.075 mg/ml were used in 25 mM Tris-HCl, 20 mM DTT, and 125 mM NaCl (pH 7.5). Prior to the $FUS^{R/G\text{-}rich\ LCD}$ droplet formation, the $His_6$-MBP tag was cleaved off by TEV protease, as described previously[1]. The RNA concentrations were chosen according to the peak position of the turbidity plots for different RNAs. The diffusion coefficient for $FUS^{R/G\text{-}rich\ LCD}$ within the condensates was calculated in a similar manner as described in our earlier work[1].

Fluorescence micrographs and FRAP data of the cluster phase were recorded using the same microscope. The cluster phase samples were made using 2.0 mg/ml peptide and 10 mg/ml RNA (past the charge inversion point) in 10 mM Tris-HCl buffer, 20 mM DTT (pH 7.5) with variable concentrations of NaCl as described in the text and/or figure legends.

*Electrophoretic mobility and size measurements:* Electrophoretic mobility of peptide-RNA droplets was measured with a dynamic light scattering (DLS) setup (ZetasizerNano ZS; Malvern Instruments Ltd.) using M3-PALS (Phase Analysis Light Scattering) technique[3]. Samples were prepared at 100 µM peptide concentration (0.22 mg/ml for $[KGKGG]_5$ and 0.24 mg/ml for $[RGRGG]_5$) and titrated against corresponding RNAs. Samples were prepared using the same buffer as the solution turbidity measurements. Each sample was incubated for at least 2 minutes. Sizes of the peptide-RNA complexes were measured using the same instrument following a similar protocol for the sample preparation.

*Partition coefficient measurements:* Peptide-RNA mixtures were prepared at the desired concentration and composition and subsequently injected into a tween 20-coated (20% v/v) 25 mm x 75 mm x 0.1 mm single chamber custom-made flow-cell. Samples were made at 0.24 mg/ml (100 μM) $[RGRGG]_5$ peptide concentration and variable RNA concentration in 25 mM Tris-HCl buffer, 25 mM NaCl, 20 mM DTT (pH 7.5). Confocal imaging was performed using a laser scanning confocal fluorescence microscope (LUMICKS™ C-trap, 60x water-immersion objective). Images were analyzed using the Fiji software[7]. For samples in the single phase region, the partition was taken to be 1.0. For the phase separated sample, partition coefficient ($k$) was calculated using the following equation:

$$k = \frac{mean\ intensity\ per\ unit\ area\ of\ droplets}{mean\ intensity\ per\ unit\ area\ of\ the\ dilute\ phase}$$

The partition coefficient estimation was carried out for several droplets per sample for statistical accuracy using Microsoft Excel. Intensity profile plots were generated using OriginPro software.

**Supplementary Note 1**

To quantitatively describe the effect of short-range attraction on the reentrant liquid condensation of peptide-RNA complexes, we first consider the charge inversion model[8]. Charge inversion occurs due to strong ion-ion correlations[9], where the multivalent polycations condense on the surface of negatively charged RNA chains. During the condensation process, the complexation of polycations with RNA happens until the total charge of the polycation-RNA complex is neutralized, *i.e.* adequate number of polycations are bound to RNA in a manner that the total positive charge is equal to the total negative charge on the complex. However, additional binding of polycations to this neutral complex may occur even after the charge neutral point, leading to accumulation of excess positive charge on the complex[10, 11]. This results in an inversion of the effective charge on the RNA chains. The energy of RNP-RNA interaction is mathematically derived from solving non-linear Poisson-Boltzmann equation and is given by[12]

$$\frac{\epsilon}{kT} = \beta \ln^2\left(\frac{C}{C_0}\right) \tag{2}$$

Where $\beta$ depends on the valency of polyions and the conventional Manning's parameter ($\zeta = \frac{l_b}{b}$). where $l_b = \frac{e^2}{\epsilon k_b T}$ is the Bjerrum length and b is the size of the polyanion monomer. As we have shown in our previous work[3], this potential, which only considers long-range ionic interactions, is sufficient to reproduce the reentrant phase transition behavior of the polycation-RNA mixtures.

Next, we consider possible short-range attraction amongst RNA and peptide chains, which alters the simplified form of the charge inversion potential. We argue that when the system has short-range attractions such as cation-$\pi$ interactions, they do not play a significant role in the thermodynamics of a dilute homogeneous mixture as compared to the long-range ionic interactions. This is simply because the average inter-particle distance ($r$) is far greater than the range of attraction ($\sim\frac{1}{r^3}\ to\ \frac{1}{r^6}$). On the contrary, long-range forces such as the screened coulomb attraction ($\sim \frac{1}{r}e^{-r/\lambda}$; λ = Debye screening length) are strong enough to drive the RNA-polycation complexation in a dilute mixture[13, 14]. This is evident from the phase separation of the polycationic peptides involved in this work with polyphosphate, which lacks RNA bases and interacts primarily via electrostatic forces (Supplementary Fig. S9). Upon complexation, a condensate is formed. The concentration of RNA and polycation is manifold higher within the condensate than the dilute mixture, and hence, the average inter-chain distance is significantly reduced. Because of this reduction in the mean intermolecular separation, short-range forces are now expected to contribute to the interaction energy more significantly than in the dilute mixture. Therefore, we represent the energy of the short-range attraction as a soft step function that increases cooperatively at the condensation boundary (Fig. 2a). The cooperative rise of short-range forces is achieved using a Gaussian Heaviside function that is described as

$$\frac{\epsilon_{short-range}}{kT} = \alpha\, \theta(x - x_c) \exp - \left(\frac{x - x_c}{\gamma}\right)^2 \tag{3}$$

Where $x_c$ is the condensation threshold, $\alpha$ and $\gamma$ are parameters that define the height and steepness of the step function, respectively. $\alpha$, the magnitude of the short-range contribution, is related to the absolute concentration of molecules inside the condensate. $\alpha$ can be estimated using any appropriate short-range potential such as Van Der Waals, Asakura-Oosawa potential, or sequence specific cation-π and/or π-π interactions. $\gamma$ describes how the concentrations of polycation and RNA change in the condensed phase along the phase boundary curve.

We next constructed a mixed potential by adding charge inversion potential and short-range attraction from equation (3). The total interaction potential is given by:

$$\frac{\epsilon_{total}}{kT} = \beta \ln^2\left(\frac{C}{C_0}\right) - \alpha\, \theta(x - x_c) \exp - \left(\frac{x - x_c}{\gamma}\right)^2 \tag{4}$$

Using this mixed potential, we performed numerical simulations of the system's phase behavior as a function of variable *strengths* of the short-range attraction (Fig. 2a&b). Our simulations reproduced the trends observed in the experimental phase boundary curves (Figs. 1b-d; 2c&d).

One important assumption of our model is that the short-range attraction remains effective after decondensation (*i.e.,* in the reentrant homogeneous phase). This is justified by our experimental observation that the RNA and the polycationic peptides form large complexes that are ≥ 200 nm in diameter (Supplementary Fig. S2) even in the reentrant phase, suggesting that the complexes are stabilized by short-range attraction against a long-range coulomb interaction.

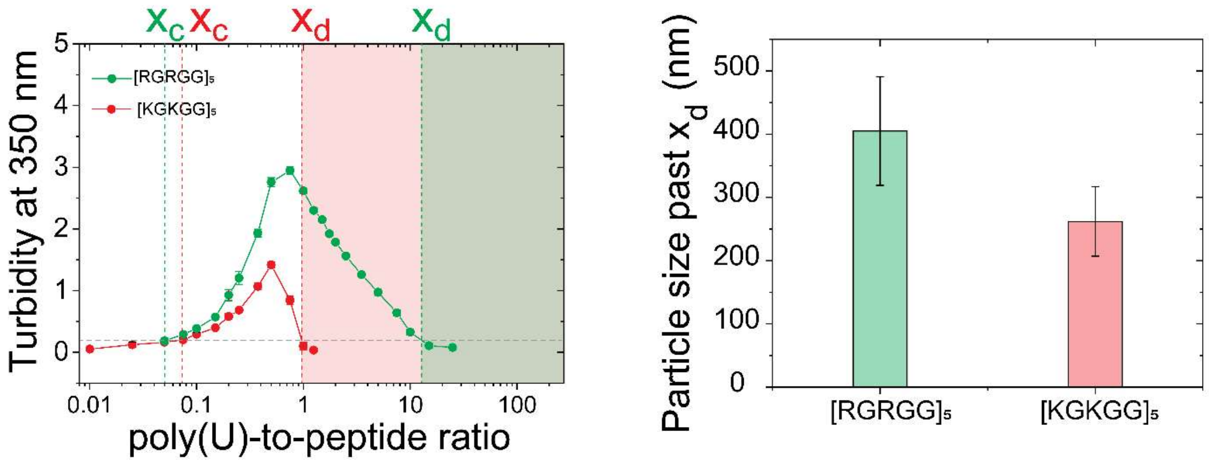


**Figure-S2 :** Particle sizes in the reentrant phase for $[RGRGG]_5$ and $[KGKGG]_5$. Phase boundary curves for $[KGKGG]_5$ and $[RGRGG]_5$ with poly(U) ***(left)***. Corresponding particle sizes as measured by dynamic light scattering (DLS) at 90° angle in the reentrant phase (past the decondensation threshold $x_d$) for the two systems ***(right)***.

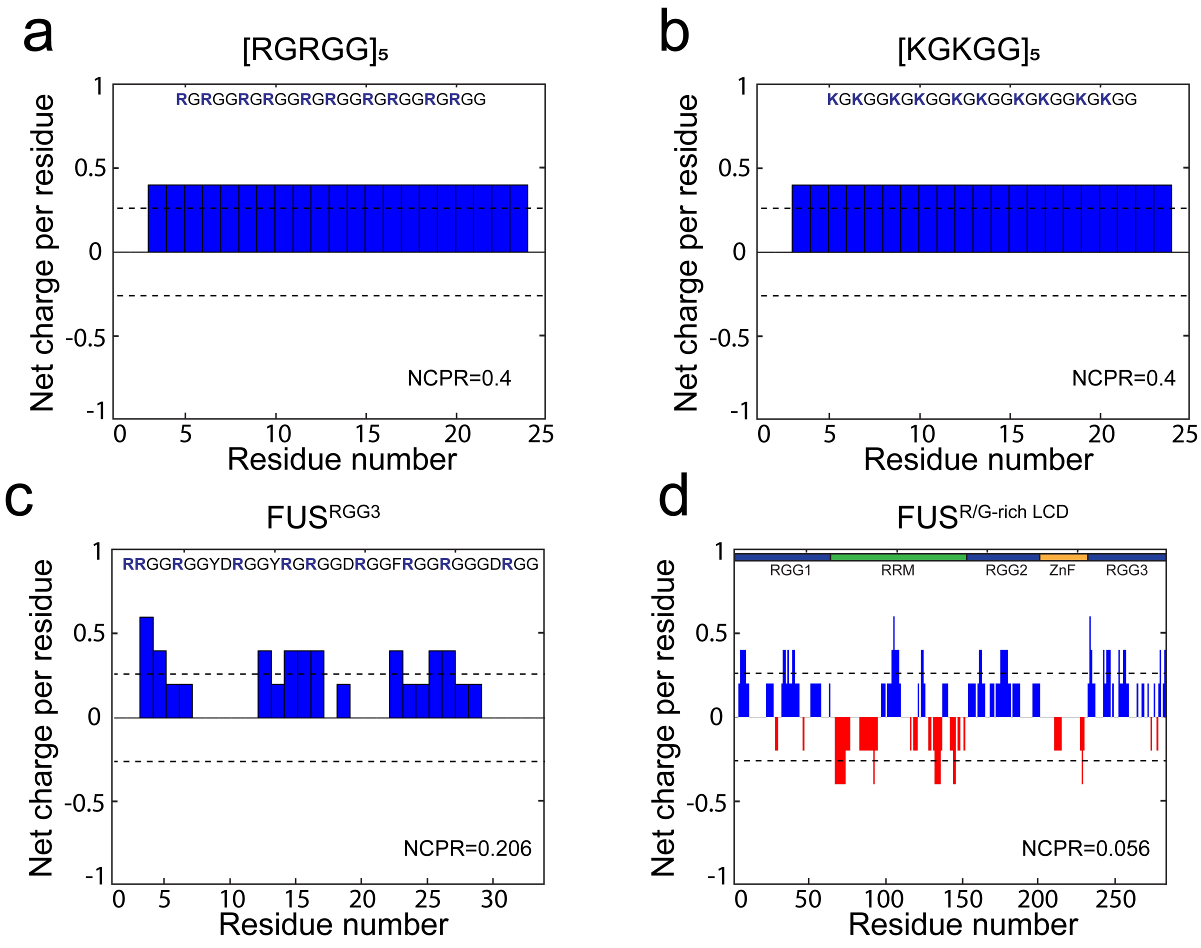


**Figure-S3 :** Net charge per residue (NCPR) distribution for **(a)** $[RGRGG]_5$, **(b)** $[KGKGG]_5$, **(c)** $FUS^{RGG3}$, and **(d)** $FUS^{R/G\text{-rich LCD}}$. NCPR was estimated using the CIDER algorithm[15].

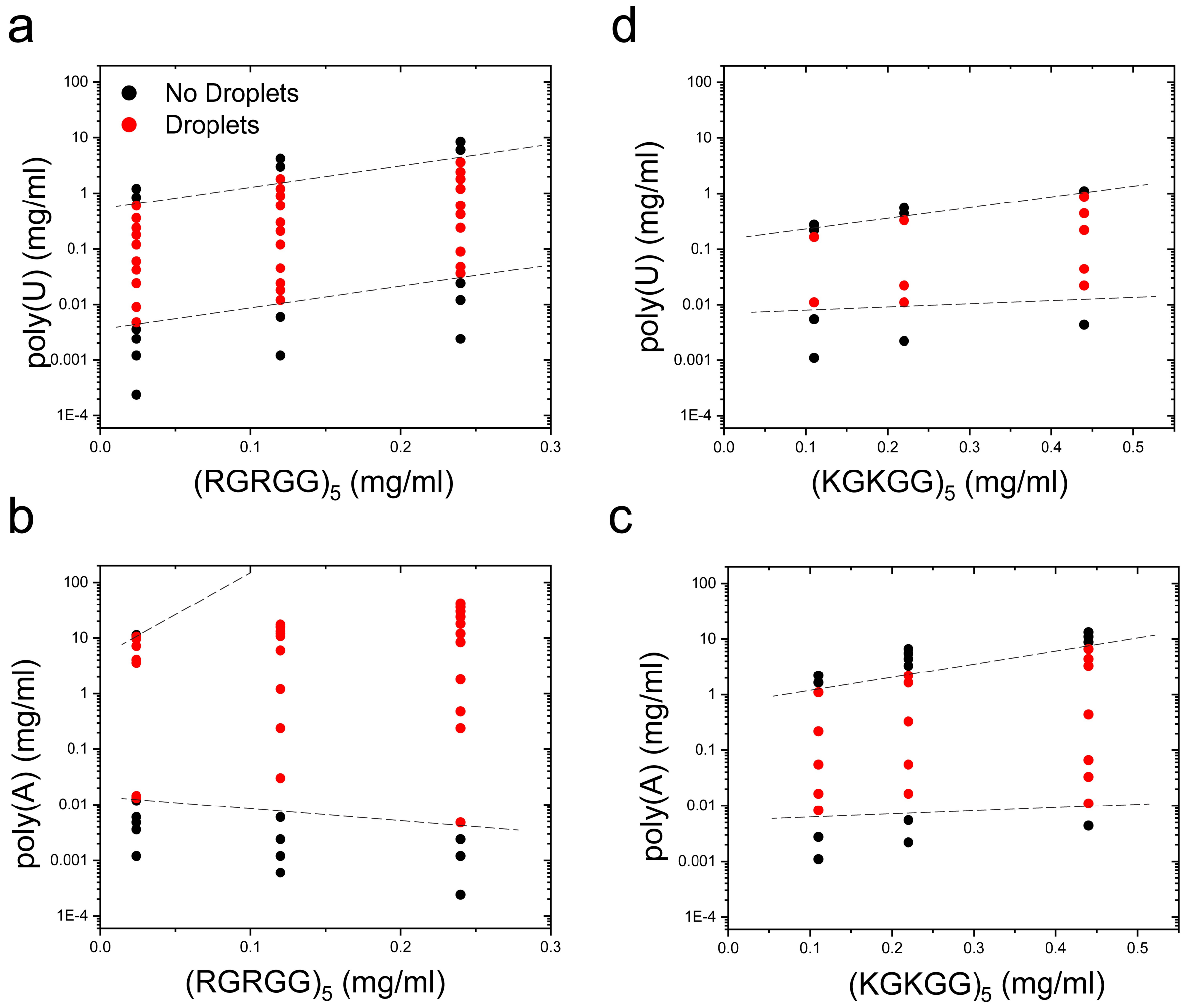


**Figure-S4 :** Phase diagrams of **(a)** $[RGRGG]_5$ with poly(U), **(b)** $[RGRGG]_5$ with poly(A), **(c)** $[KGKGG]_5$ with poly(A), and **(d)** $[KGKGG]_5$ with poly(U) RNA. Red and black points represent two phase regions and one phase regions, respectively. Dotted lines represent phase boundaries. Buffer used: 25 mM Tris-HCl, 20 mM DTT (pH 7.5).

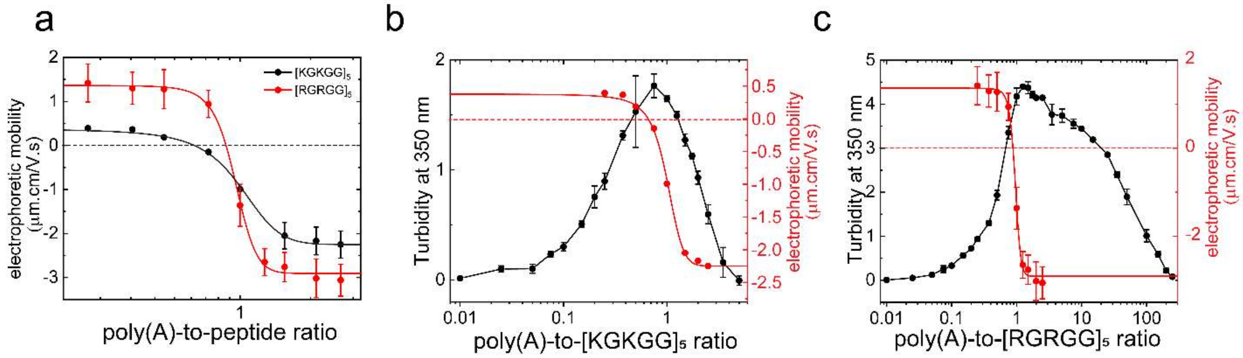


**Figure-S5**: **(a)** Electrophoretic mobility of poly(A)-[RGRGG]$_5$ and poly(A)-[KGKGG]$_5$ condensates as a function of poly(A)-to-peptide ratio. Solid lines are shown as simple guide to the eye. **(b)** Overlay plot of turbidity at 350 nm (left axis, black) and electrophoretic mobility (right axis, red) for poly(A)-[KGKGG]$_5$ mixture. **(c)** Overlay plot of turbidity at 350 nm (left axis, black) and electrophoretic mobility (right axis, red) for poly(A)-[RGRGG]$_5$ mixtures. (buffer: 25 mM Tris-HCl, pH 7.5, containing 20 mM DTT).

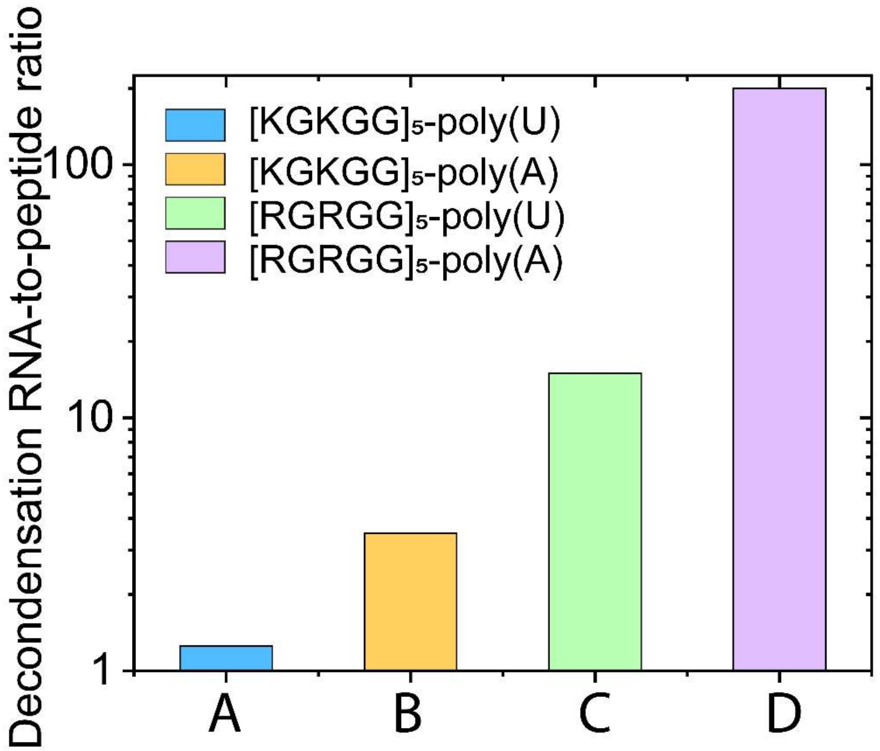


**Figure-S6:** Decondensation boundaries corresponding to the emergence of a reentrant phase (from the macroscopic turbidity assay data presented in **Figure. 1** in the maintext). **(A)** poly(U)-[KGKGG]$_5$, **(B)** poly(A)-[KGKGG]$_5$, **(C)** poly(U)-[RGRGG]$_5$, **(D)** poly(A)-[RGRGG]$_5$.

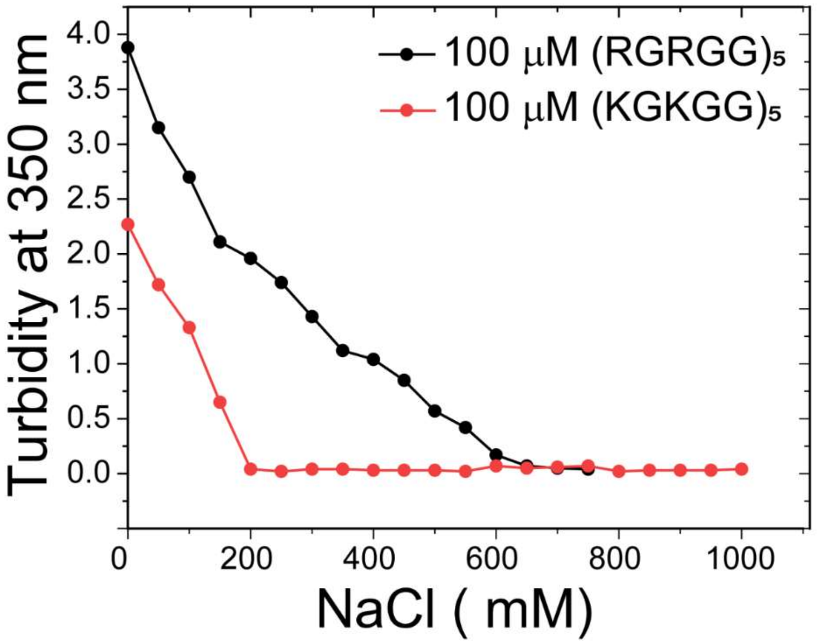


**Figure-S7: Effects of ionic strength on the stability of peptide-RNA droplets.** Solution turbidity at 350 nm for independently prepared mixtures of $[RGRGG]_5$ (100 μM or 0.24 mg/ml; black) and $[KGKGG]_5$ (100 μM or 0.22 mg/ml; red) with poly(U) RNA at 1:1 ratio as a function of salt concentration. Buffer condition: 25 mM Tris-HCl, 20 mM DTT (pH 7.5) with variable NaCl concentrations as indicated in the plot.

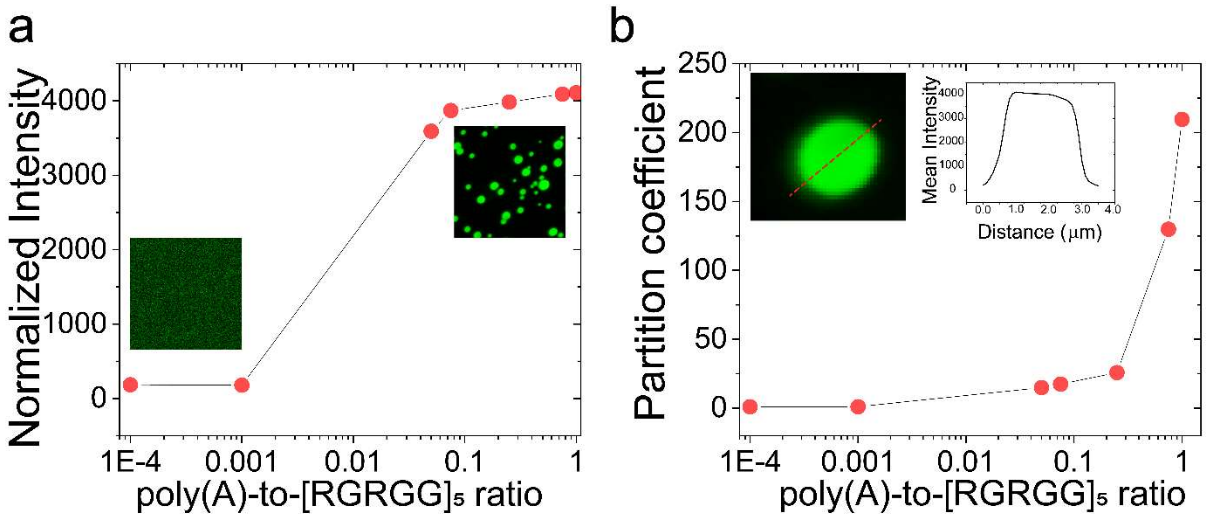


**Figure-S8:** Intensity and partition coefficient analysis during the transition from a mixed phase (regime I) to the condensed phase (regime II) for $[RGRGG]_5$-poly(A) mixture at a peptide concentration of 0.24 mg/ml. (buffer: 25 mM Tris-HCl, pH 7.5, 20 mM DTT, 25 mM NaCl).

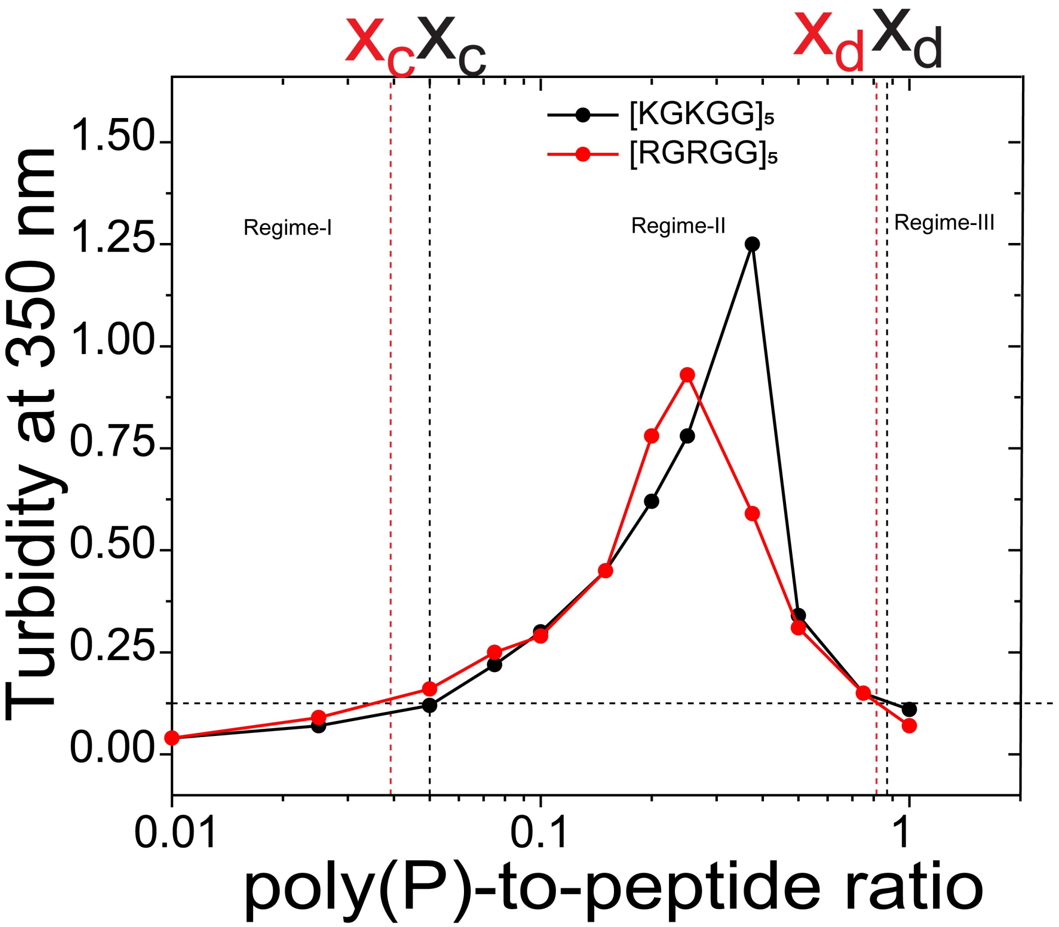


**Figure-S9:** Solution turbidity at 350 nm for $[RGRGG]_5$ and $[KGKGG]_5$ as a function of polyphosphate-to-peptide ratio. Peptide concentration: 100 μM, buffer: 25 mM Tris-HCl, 20 mM DTT (pH 7.5).

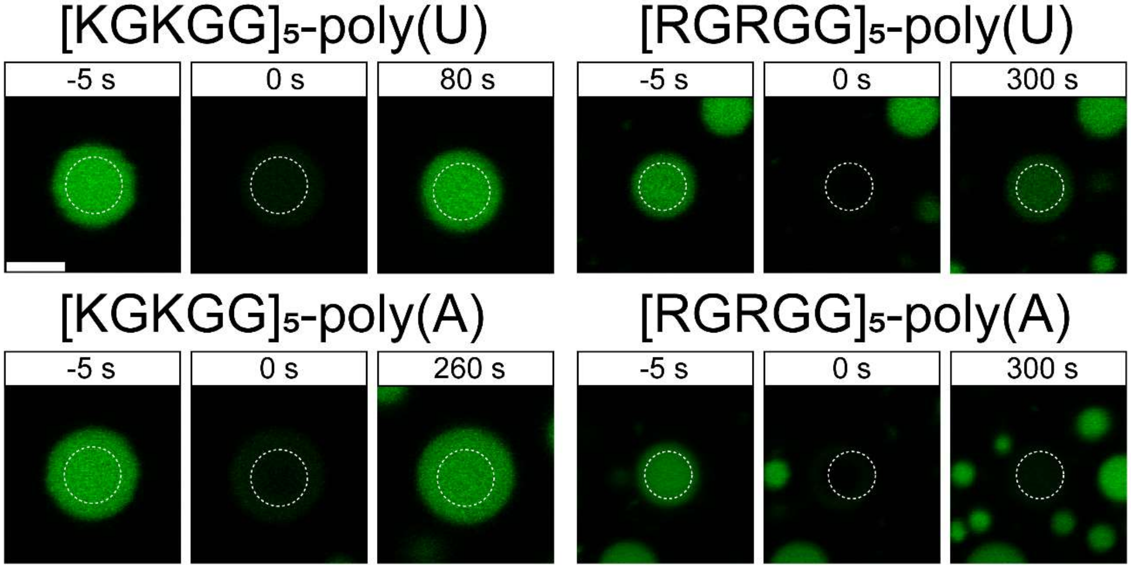


**Figure-S10:** Time-lapse FRAP images for various peptide-RNA systems. Negative time points indicate pre-bleaching times (bleaching occurs at t=0 s). Scale bar represents 5 µm. The FRAP plots are shown in Fig. 3 in the maintext. All samples for FRAP measurements were prepared at 1.0 mg/ml peptide concentration, 0.75 mg/ml RNA concentration (corresponding to the peak in their turbidity plots shown in Fig. 1b-d) and a buffer (25 mM Tris-HCl, 20 mM DTT, 25 mM NaCl (pH 7.5)).

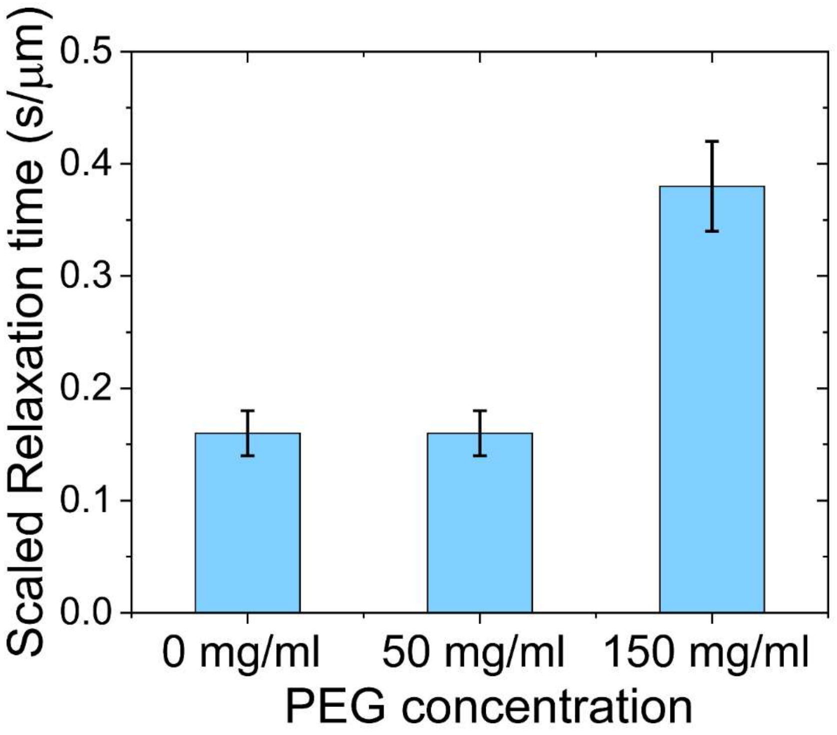


**Figure-S11**: Fusion relaxation time as a function of crowder concentration for $FUS^{RGG3}$-poly(U) condensates. Samples were prepared using a peptide concentration of 0.33 mg/ml at a poly(U)-to-$FUS^{RGG3}$ ratio of 0.75 (buffer: 25 mM Tris-HCl, 150 mM NaCl (pH 7.5)).

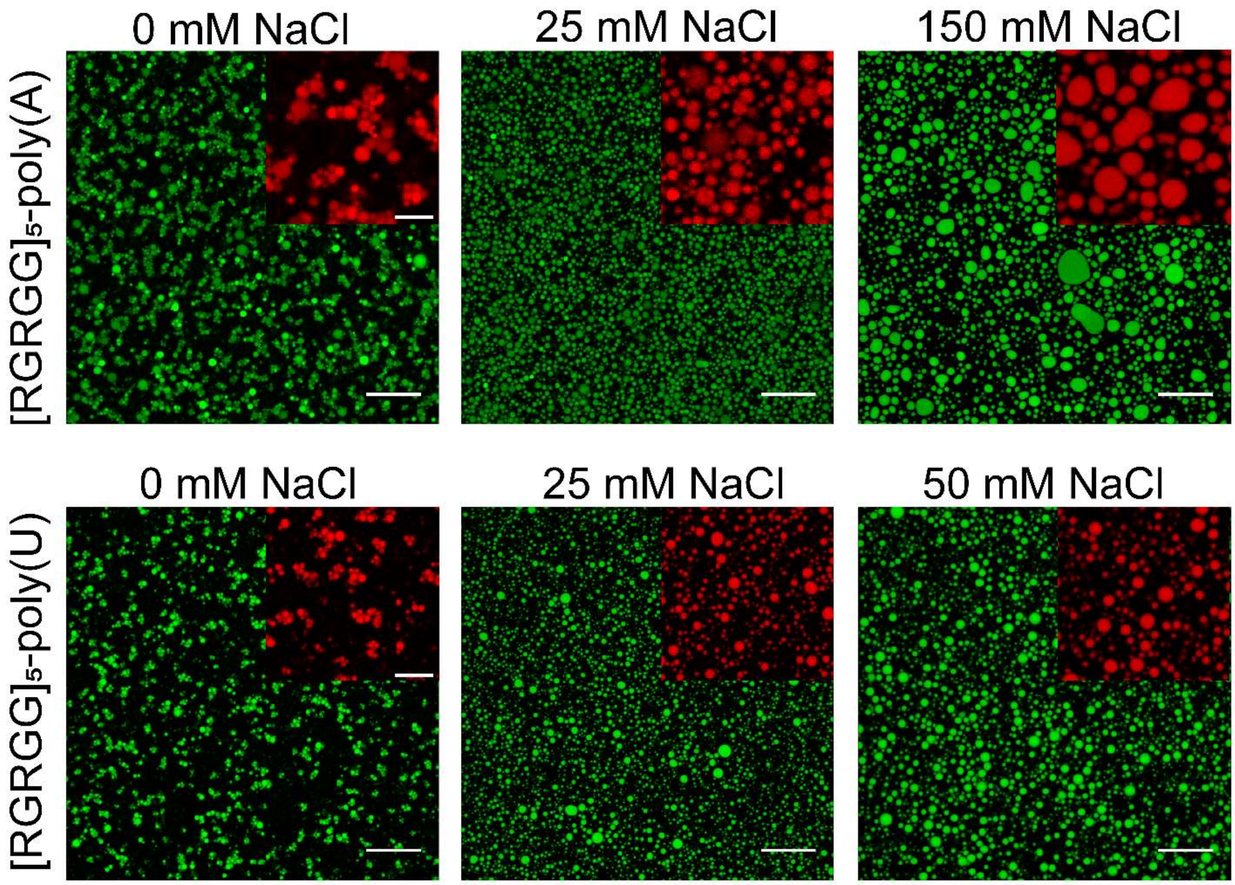


**Figure S12:** Droplet clusters transition to a bulk liquid phase with increasing salt concentration. Shown here are droplet clusters formed by poly(A)-$[RGRGG]_5$ mixture (upper panel) and poly(U)-$[RGRGG]_5$ mixture (lower panel). Poly(A) samples were prepared at 2.0 mg/ml $[RGRGG]_5$, 10 mg/ml poly(A) in 10 mM Tris-HCl, 20 mM DTT (pH 7.5) buffer and images are shown for 0 mM, 25 mM and 150 mM NaCl concentrations (from left to right). Poly(U) samples were preparaed at similar peptide/RNA concentrations in 5.0 mM Tris-HCl buffer (pH 7.5), 20 mM DTT and images are shown for 0 mM, 25 mM and 50 mM salt (left to right). The *insets* are zoomed in view with pseudocolored red for better clarity. Scale bar represents 20 µm (green images) and 10 µm (red images), and are shown only for the first image in each sequence. The rest of the images in a given sequence follow the same scale bar as the initial image.

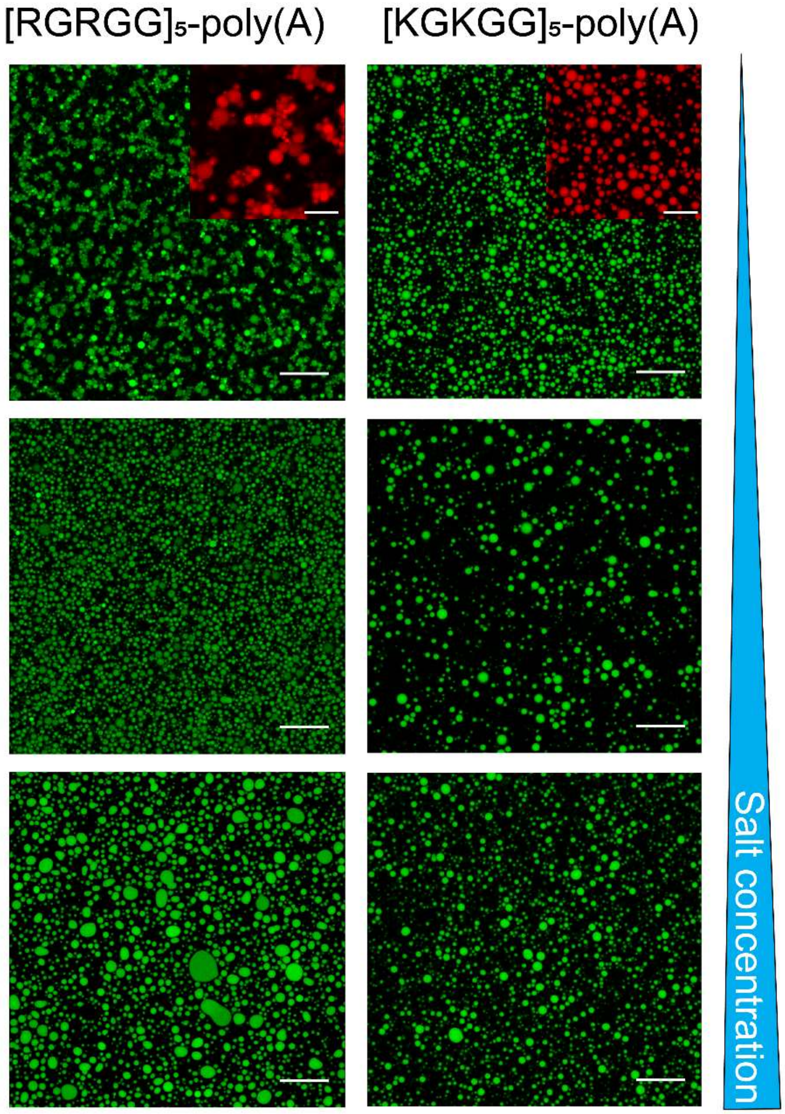


**Figure-S13:** Fluorescence micrographs showing droplet clusters (or a lack thereof) as a function of salt. **(Left panel)** poly(A)-[RGRGG]$_5$ droplet clusters at low salt buffer (10 mM Tris-HCl, pH 7.5, 20 mM DTT) transitions to droplet suspensions upon increase in buffer ionic strength. **(Right panel)** poly(A)-[KGKGG]$_5$ mixtures displaying droplet suspensions at low salt buffer (10 mM Tris-HCl, pH 7.5, 20 mM DTT) that grow with increase in buffer ionic strength. Both samples were prepared using 2.0 mg/ml peptide and 8.0 mg/ml poly(A). Salt concentrations used in these experiments are 0, 25, and 50 mM NaCl for each row from top to bottom. Scale bar is 20 µm for green images and 10 µm for red images. The *insets* are zoomed in view pseudocolored red for better clarity.

**Supplementary Table-1**

| pH | Sequence net charge | | | |
|---|---|---|---|---|
| | $[RGRGG]_5$ | $[KGKGG]_5$ | $FUS^{R/G\text{-rich LCD}}$ | $FUS^{RGG3}$ |
| 6.5 | 10 | 10 | 17.7 | 7 |
| 7 | 9.9 | 9.9 | 16.7 | 6.9 |
| **7.5** | **9.8** | **9.7** | **16** | **6.8** |
| 8 | 9.5 | 9.4 | 15.4 | 6.5 |
| 8.5 | 9.2 | 8.9 | 14.7 | 6.2 |

**Supplementary Table-1:** Estimated net charge for $[RGRGG]_5$, $[KGKGG]_5$, $FUS^{R/G\text{-rich LCD}}$, and $FUS^{RGG3}$. The corresponding sequences are shown in Supplementary Figure S3. The net charge is estimated using protein calculator v3.4 developed by *C.D Putnam at the Scripps research institute* (http://protcalc.sourceforge.net/).

**Supplementary Movie Legends:**

**Supplementary Movies 1-4:** Trap-mediated coalescence of suspended droplets of **(1)** $[RGRGG]_5$-poly(A), **(2)** $[RGRGG]_5$-poly(U), **(3)** $[KGKGG]_5$-poly(U), and **(4)** $[KGKGG]_5$-poly(A). Scale bar represents 1 µm for **(1)** and 2 µm for **(2-4)**.

**Supplementary Movies 5-6:** Trap-mediated coalescence of suspended droplets of **(5)** $FUS^{R/G\text{-}rich\ LCD}$-poly(A) **(6)** $FUS^{R/G\text{-}rich\ LCD}$-poly(U). Scale bar represents 2 µm.

**Supplementary Movie 7:** Temporal dynamics of $[RGRGG]_5$ -poly(A) droplet clusters. Scale bar represents 10 µm.

**Supplementary Movies 8-9:** FRAP movies of clusters formed by **(8)** $[RGRGG]_5$-poly(A) and **(9)** $[RGRGG]_5$-poly(U). Scale bar represents 10 µm.
